# Tryptophan Metabolites And Their Predicted Microbial Sources In Fecal Samples Of Healthy Individuals

**DOI:** 10.1101/2023.12.20.572622

**Authors:** Cynthia L. Chappell, Kristi L. Hoffman, Philip L. Lorenzi, Lin Tan, Joseph F. Petrosino, Richard A. Gibbs, Donna M. Muzny, Harsha Doddapaneni, Matthew C. Ross, Vipin K. Menon, Anil Surathu, Sara J. Javornik Cregeen, Anaid G. Reyes, Pablo C. Okhuysen

## Abstract

Gut microbiota produce tryptophan metabolites (TMs) important to homeostasis. However, measuring TM levels in stool and determining their microbial sources can be difficult. Here, we measured TMs from the indole pathway in fecal samples from 21 healthy adults with the goal to: 1) determine fecal TM concentrations in healthy individuals; 2) link TM levels to bacterial abundance using 16S and whole genome shotgun (WGS) sequencing data; and 3) predict likely bacterial sources of TM production. Within our samples, we identified 151 genera (16S) and 592 bacterial species (WGS). Eight TMs were found in ≥17 fecal samples, including four in all persons. To our knowledge, we are the first to report fecal levels for indole-3-lactate, indole-3-propionate, and 3-indoleacrylate levels in healthy persons. Overall, indole, indole-3-acetate (IAA), and skatole accounted for 86% of the eight TMs measured. Significant correlations were found between seven TMs and 29 bacterial species. Predicted multiple TM sources support the notion of a complex network of TM production and regulation. Further, the data suggest key roles for *Collinsella aerofaciens* and IAA, a metabolite reported to maintain intestinal homeostasis through enhanced barrier integrity and anti-inflammatory/antioxidant activities. These findings extend our understanding of TMs and their relationship to the microbial species that act as effectors and/or regulators in the healthy intestine and may lead to novel strategies designed to manipulate tryptophan metabolism to prevent disease and/or restore health to the dysbiotic gut.

**IMPORTANCE:** Tryptophan metabolites (TMs) of bacterial origin are increasingly recognized as important signaling molecules among gut microbiota and with the host. However, few reports exist for fecal TM levels in healthy humans, and reported levels vary widely. Further, the specific bacterial species producing TMs and the combinations of fecal TMs in healthy individuals are not well known. Our research combines 16S and whole genome shotgun sequencing of gut bacteria with a sensitive method (LC/MS) for measuring TMs and a reported method to predict which species are likely TM contributors. To our knowledge, this combination of analyses has not been reported elsewhere and will add significantly to the existing literature. Understanding TM levels and their sources in the healthy intestine are fundamental to elucidating how TMs contribute to maintaining homeostasis. Such knowledge of gut microbiota and their metabolic products will inform novel strategies to maintain intestinal health and prevent or treat dysbioses.

## INTRODUCTION

The intestinal microbial community is shaped by diet, environment, age, ethnicity, health status, and the host immune response, among others (1–3). Attempts to define a “healthy” microbiome have led to the understanding that necessary functions can be fulfilled by a variety of species (4), a characteristic important in adaptation to new environments or conditions. This understanding extends to the molecular signals, including tryptophan metabolites (TMs), produced and secreted by commensal bacteria that contribute to gut homeostasis.

Resident gut microbiota are an important and often sole source of indole metabolites as evidenced by the lack of these metabolites in gnotobiotic (5) and antibiotic-treated mice (6). These metabolites influence a variety of functions within the microbial community by affecting biofilm formation, antimicrobial activity, drug resistance, and expression of virulence factors (reviewed in 7-8). Likewise, TMs (Figure 1) have profound effects on the host mucosal epithelium, modulation of the immune response, gut hormone secretion, and various other intestinal functions (reviewed in 7-11). Microbial-derived TMs have also been implicated as effectors in extraintestinal tissues, such as in the gut-brain axis (reviewed in 12; 13-14), and many studies to date have documented altered TM concentrations in multiple intestinal diseases and conditions (reviewed in 7,9). In addition, a body of literature has implicated microbial-derived TMs as effectors in extraintestinal tissues, such as in the gut-brain axis (reviewed in 12; 13-14).

**Figure 1.**
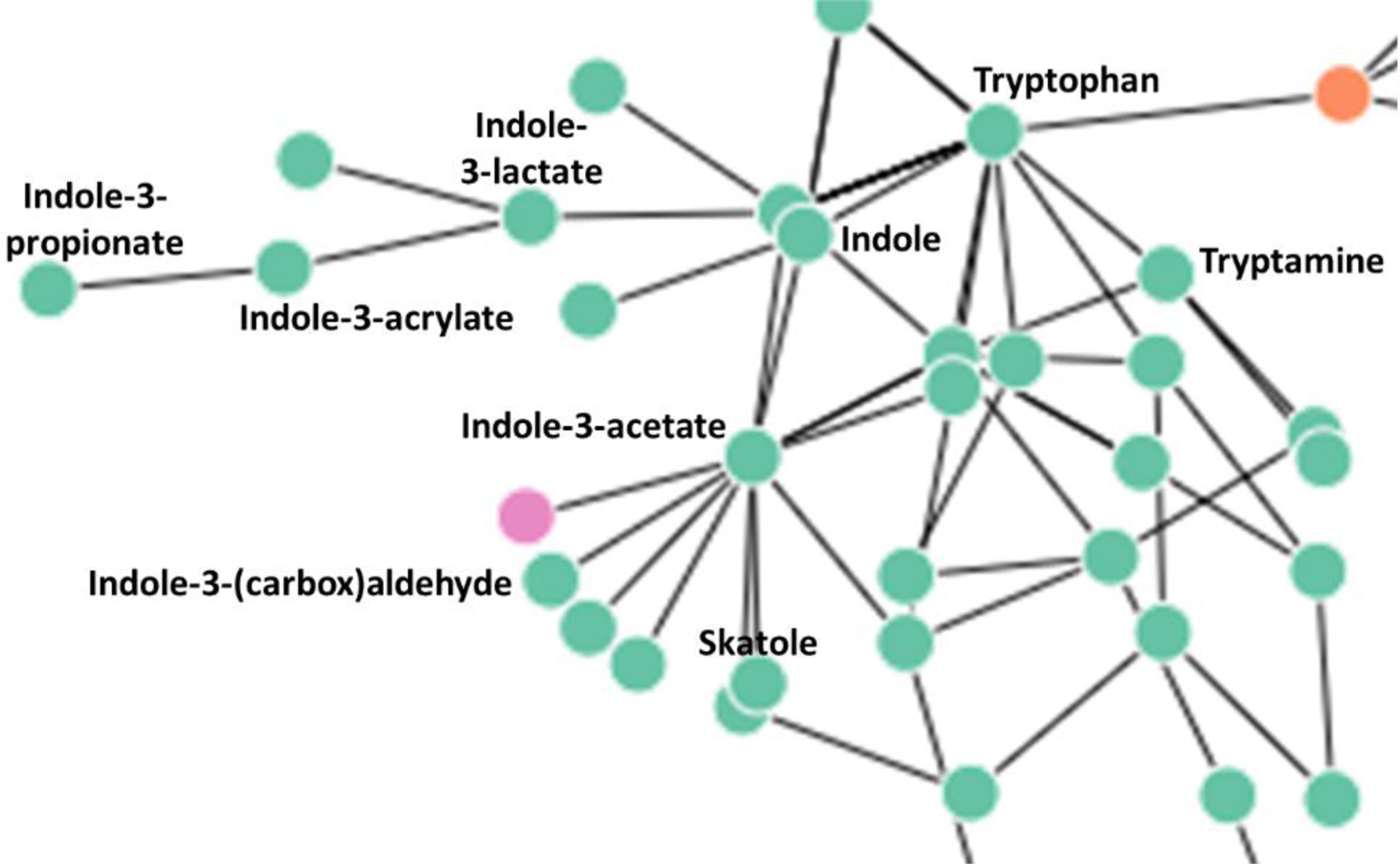
Canonical pathways for tryptophan metabolism in bacteria. The indole pathway is adapted from Lu et al., 2022; www.TrpNet.ca. Metabolites included in the present study are indicated. Orange and truncated lines indicate the metabolic connections to host specific pathways; pink color indicates an unknown enzymatic reaction.

Despite a growing recognition of their importance, fecal TMs and their microbial sources in healthy humans are understudied. Previous studies, although few in number, identified up to five TMs in human stool, while two other metabolites, IPA and IA, were detected in rodent feces (15; reviewed in 16; Table S1). Other reports of these metabolites have been in the context of gastrointestinal disease or inflammation and reference values from healthy controls are lacking.

This relative dearth of information is, in part, due to the difficulty of quantitating most of these metabolites. Only three TMs (indole,IND; skatole, SK; and indole acetic acid, IAA) can be measured using spectrophotometric assays. The Kovacs test, first described in 1928 (17), detects indole and skatole together compared to hydroxylamine assay, which specifically detects indole (18). IAA can be detected using a modification (19) of the Salkowski assay (20), but the method is not specific and therefore not useful for complex biological samples. Thus, TM detection and quantitation currently relies on gas or liquid chromatography followed by mass spectrometry, a process that requires sophisticated instrumentation and expertise. Thus, the actual number and identity of bacterial species that contribute to the production of fecal TMs are not fully known.

In the past, identification of TM-producing bacteria relied on detecting the metabolites in culture supernatants (reviewed in 7,21). *In silico* studies expanded identification but, of necessity, relied on bacteria for which a complete genome sequence existed (22). Together, these studies revealed that not all species within a bacterial genus have the capacity to produce TMs. The recent analytical tool TrpNet (Trpnet.ca)goes even further using a Bayesian logistic regression model to assign a metabolite production probability (MPP) for all known TMs associated with mouse and human gut bacteria (at different taxonomic levels) (23). MPPs can then be applied to a taxon’s relative abundance to predict likely sources of a specific metabolite in individual fecal samples. Past studies have also related TM levels in body fluids to qualitative and/or quantitative changes in the gut microbiome (reviewed in 7-8).

The overall purpose of this study was to add and/or extend information regarding TM concentrations in the healthy human gut and to identify their potential microbial sources. Specifically, our objectives were to: 1) obtain 16S and whole genome sequences from 21 healthy adults; 2) determine the concentrations of ten (10) biologically active tryptophan metabolites in fecal samples; 3) identify potentially important taxa associated with TM production; 4) use the new web-based tool TrpNet to predict specific microbial sources of TMs and compare these species to observed TM concentrations. These findings will add significantly to the literature regarding the role of gut bacteria and their metabolites in human health and disease. Understanding the contribution of specific bacterial species and their TMs will further support a rational approach to drug and/or dietary interventions for preventing and/or treating dysbioses.

## MATERIALS AND METHODS

### Study population and experimental approach

Stool samples used in the present study were collected in 1993-1995 from healthy adults as part of a *Cryptosporidium* challenge study approved by the Committee for the Protection of Human Subjects at the University of Texas Health Science Center in Houston (#HSC-MS-02-105). Volunteers underwent a series of examinations and tests to verify that all were healthy with no underlying conditions or gastrointestinal illnesses within the previous six months. Demographic information was recorded at the initial enrollment in the study. Baseline stool samples were collected after enrolment and kept on ice until delivery to the laboratory (within 24 hours). Samples were immediately frozen and stored at −80C but later thawed, aliquoted, and refrozen for batch processing. One sample had additionally been used in an earlier study (18) where it was partially thawed until a portion could be scrapped from the top, and then the remainder was immediately refrozen at −80C.

### DNA extraction, library preparation, and sequencing

Stool aliquots were delivered on dry ice to the Alkek Center for Metagenomics and Microbiome Research at Baylor College of Medicine, Houston, TX (J. Petrosino) for processing and sequencing. Total genomic DNA was extracted using the Qiagen DNeasy PowerSoil Pro HT Kit on the Qiagen QIAcube HT liquid handling platform and quantified using a Picogreen assay. Libraries for 16S rRNA gene sequencing were prepared as previously described (24) and sequenced on the Illumina MiSeq platform using the 2×250bp paired-end protocol.

WGS libraries were constructed beginning by first shearing extracted DNA (20 ng to 500 ng) was sheared into fragments of approximately 200-600 bp on a Covaris E220 system. Shearing was followed by double-size selection purification using AMPure XP beads. DNA end repair, 3’-adenylation, ligation to Illumina multiplexing PE adaptors, and ligation-mediated PCR (LM-PCR) were all completed using automated processes. A set of 96 Unique dual index barcodes (Illumina TruSeq UD Indexes, # 20022370) were utilized to barcode samples. Libraries were amplified for 13 PCR cycles in 50 μl reactions containing 150 pmol of P1.1 (5’-AATGATACGGCGACCACCGAGA) and P3 (5’-CAAGCAGAAGACGGCATACGAGA) primer and Kapa HiFi HotStart Library Amplification kit (Cat# kk2612, Roche Sequencing and Life Science). Library quantification and size estimation were performed using a Fragment Analyzer (Agilent Technologies, Inc.) electrophoresis system and the final library product was selected at 470-580bp. Prepared libraries were then pooled and sequenced according to the experimental plan and handed off for sequencing on an Illumina NovaSeq 6000 S4 flow cell, using the 2×150 bp paired-end protocol.

### Bioinformatics processing

Raw data files in binary base call (BCL) format were converted into FASTQs and demultiplexed based on the dual-index barcodes using the Illumina ‘bcl2fastq’ software. Demultiplexed raw fastq sequences were processed using BBDuk to quality trim, remove Illumina adapters, and filter PhiX reads. Trimmed FASTQs were mapped to a combined PhiX and host reference genome database using BBMap to determine and remove host/PhiX reads. Taxonomic profiling of sequenced samples was determined using MetaPhlAn3 following the recommended workflow with a modification to the read mapping tool. Briefly, the processed FASTQ reads were mapped against the MetaPhlAn3 marker gene database (mpa_v30_CHOCOPhlAn_201901) using BBMap before processing through the MetaPhlAn3 algorithm. This created the default relative abundance and estimated counts table per kingdom per sample.

### Rarefaction

Rarefaction depths were 2272 (16S) and 1451921 estimated counts (WGS) reads (Figure S1). The combined sequencing methods identified 324 genera (classified and unclassified; 16S, 224; WGS, 197), and 97 genera identified by both methods.

Rare taxa, i.e. found in <20% of individuals, were excluded from analyses. The remaining commonly occurring taxa accounted for a median relative abundance of 97.6% (16S) and 99.9% (WGS), ranging from 80.1 - 100% and 91.5 - 100%, respectively, per sample. Likewise, WGS yielded 137 species identified in >20% of subjects, representing a median of 99.3% (range, 89.7 - 99.9%).

### Tryptophan metabolite (TM) quantitation

Indole compounds used in the study are listed along with their chemical structures (Table 1). Purified metabolites for standard curves were purchased from: TCI America, Portland OR (IA, IAA, TAM); Chem-Impex International Inc, Wood Dale, IL (ILA); Acros Organics, Waltham, MA (IND, I-Ald, IAc); Alfa Aesar, ThermoFisher Scientific, Tewksbury, MA (SK, I3S); and MP Biochemicals, LLC, Illkirch, France (IPA). Frozen fecal samples (100-200 mg) were delivered on dry ice to the Metabolomics Facility at

**Table 1.**
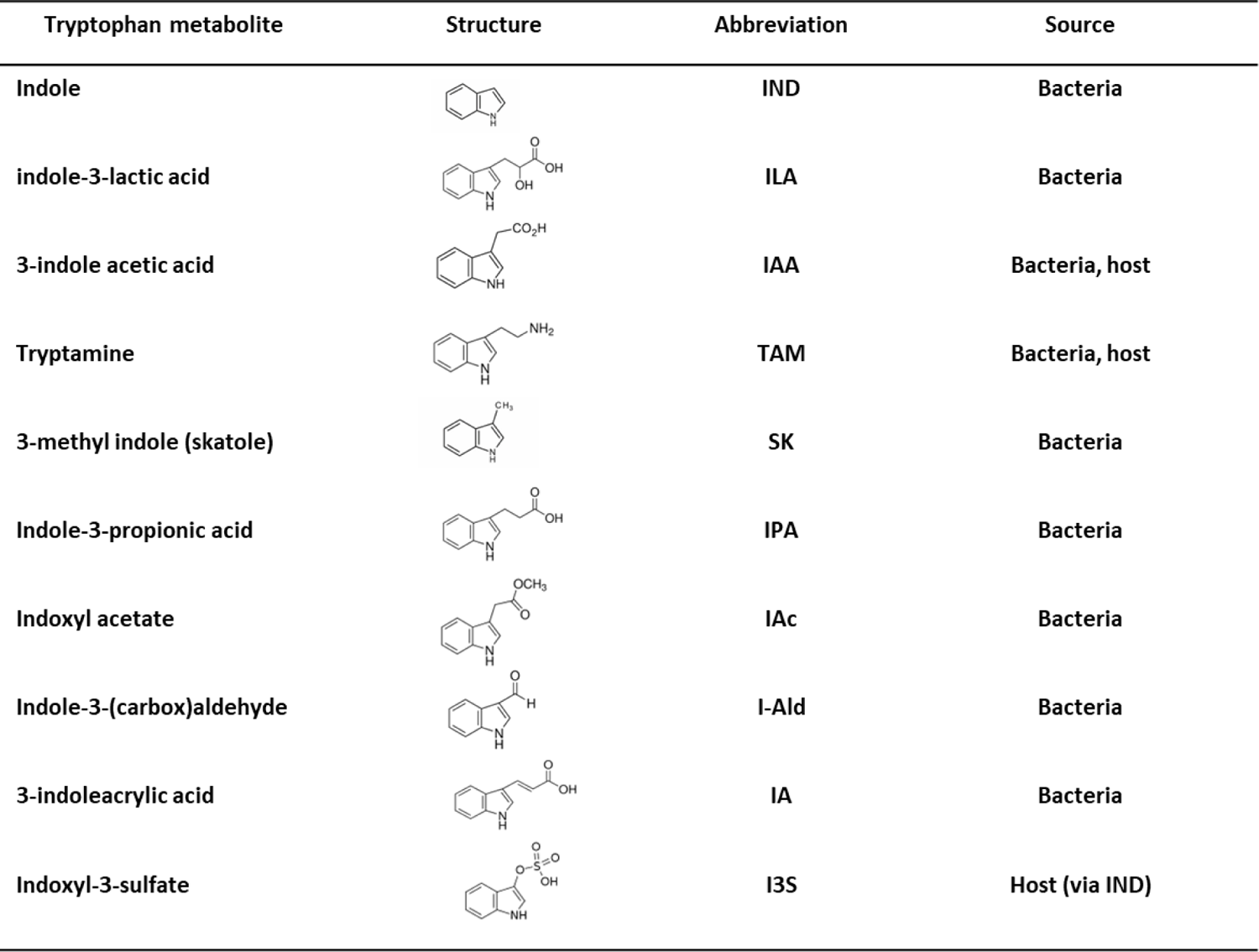
Tryptophan metabolites, their structures, abbreviations and sources.

M.D. Anderson Cancer Center (P. Lorenzi, Houston, TX) for extraction and testing. To determine the relative concentrations of tryptophan metabolites, extracts were prepared and analyzed by liquid chromatography coupled with high-resolution mass spectrometry (LC-HRMS). Samples were thawed on ice, and metabolites were extracted by adding 0.5 mL ice-cold 50/50 (v/v) methanol/acetonitrile and homogenizing with a Precellys Tissue Homogenizer. The extract was centrifuged at 17,000 *g* for 5 min at 4°C, and the supernatant was transferred to a clean tube. A subsequent round of extraction was performed by adding another 0.5 mL 0.1% formic acid in 50/50 (v/v) acetonitrile/water to the same sample, then re-homogenized. The supernatant was combined with previous step to get a total volume of 1.0 mL, followed by evaporation to dryness under nitrogen. Samples were reconstituted in 0.1 mLs of 50/50 (v/v) methanol/water, then 10 μL was injected into a Thermo Vanquish high pressure liquid chromatography (HPLC) system containing a Waters XSELECT HSS T3 2.1x 150 mm column with 2.5 µm particle size. Mobile phase A (MPA) was 0.1% formic acid in water. Mobile phase B (MPB) was 100% methanol. The flow rate was 200 µL/min (at 35°C), and the gradient conditions were: initial 5% MPB, increased to 95% MPB at 15 min, held at 95% MPB for 5 min, returned to initial conditions and equilibrated for 5 min. The total run time was 25 min. Data were acquired using a Thermo Orbitrap Fusion mass spectrometer under ESI positive and negative ionization modes at a resolution of 240,000 with full scan mode. Raw data files were imported into Thermo Trace Finder software for final analysis. The relative concentration of each compound was normalized by stool weight, and data were expressed as nmol/g. For the identified metabolites, total TM was calculated for each sample by summing the metabolites (nmol/g) present. Relative metabolite levels (%) in each sample were also calculated by using total TM level as the denominator.

### Correlation of tryptophan metabolite concentration and taxon relative abundance

The relationship between TM level and taxon abundance was examined by linear regression correlation. WGS identified a total of 197 genera (including unclassified) and 592 species in the study population with no restriction on % occupancy. Of these, 77 genera were also identified by TrpNet as “TM-producers”. An expanded analysis included all species (n=174) in these 77 genera regardless of their metabolite-producing status. Using *Agile Toolkit for Incisive Microbial Analyses* (ATIMA; https://atima.research.bcm.edu), species %RA was plotted against each of eight metabolite concentrations (nmol/g), and linear regression correlations (r^2^) and FDR-adjusted p values were obtained. A less specific, but broader approach using 16S data was also undertaken so that no potentially important taxa were overlooked. Using the same selection process, we identified 33 TM-producing genera, which were analyzed as above.

### Predicted microbial sources of tryptophan metabolites

Potential producers of one or more TMs included genera (16S) and/or species (WGS) identified by TrpNet (23, www.trpnet.ca), which also assigned metabolite production probabilities (MPPs) for each TM. Since MPPs were not available for unclassified or unnamed species, the probability for the next highest taxonomic level was used as an estimate. It should be noted that MPPs at the genus or higher taxonomic levels represent an average MPP for each species, not all of which are TM producers. A prediction score (PS), the likelihood of a taxon as a potential source of metabolite production, was calculated per fecal sample for each taxon and TM as follows: MPP (limited to >0.1 probability) X percent relative abundance (%RA). Further, to verify the predictive value of the approach, regression analysis was used to plot PS for each taxon against TM level (nmol/g) per sample, and the r^2^ and curve of best fit were determined.

### Data analysis and statistical tests

Unless otherwise indicated, taxa were included in the analysis if they were found in >20% (i.e. ≥5) of 21 individuals in the study population. Median values and non-parametric tests were used when data were not normally distributed. Statistical tests are indicated in the legends of figures and tables. Descriptive statistics and tests for significance were done using Prism (GraphPad Prism version 9.2.0 for Windows, GraphPad Software, San Diego, California USA, www.graphpad.com) or ATIMA.

Relationships with continuous variables were tested using R’s base function for linear regression models. All p-values were adjusted for multiple comparisons with Benjamini and Hochberg’s formula for the false discovery rate. Chord diagrams for correlations and prediction scores were constructed using datasmith.org.

## RESULTS

### Description of study population and experimental approach

Historical samples used in the present study consisted of a single stool sample from each of 21 healthy individuals. Mean age was 32.1 years (range 22-45) with males significantly younger (median= 28.5; Mann-Whitney statistic, p= 0.029) than females (median= 37); females constituted 61.9% of the subjects. A majority were non-Hispanic White (57.1%; 7 males, 5 females), with non-Hispanic Black (28.6%; 0 males, 6 females) and Hispanic individuals (14.3%; 1 male, 2 females) also represented (Table S2).

The study design and analytical approach are outlined (Figure 2). Fecal samples were assessed for TM concentrations by mass spectroscopy, and bacteria were sequenced. Using these datasets, four types of analyses were undertaken to: determine relationships among TM concentrations (analysis step 1); indicate which taxa influenced (or were influenced by) particular TMs (analysis step 2); generate prediction scores (PS) indicating the relative likelihood of taxa as TM sources in individual samples (analysis step 3); and lastly, test the strength of the predicted microbial sources in comparison to observed TM levels (analysis step 4).

**Figure 2.**
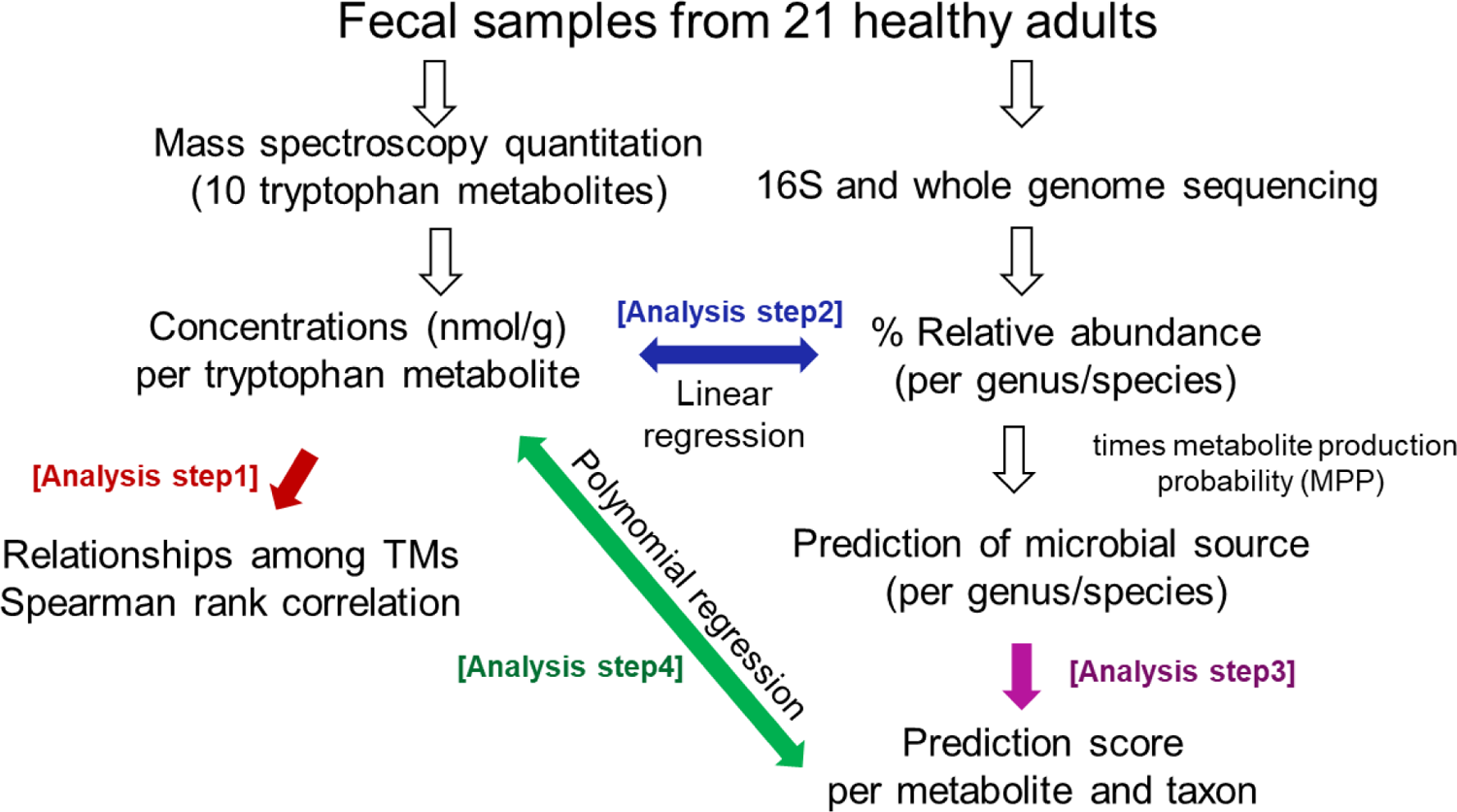
Study design and analytical approach. Analytical steps 1-4 are indicated by color.

### Tryptophan metabolite quantitation in stool samples

Eight TMs in the canonical indole pathway (Figure 1) were detected and measured by LC-MS, including six of exclusive bacterial origin (Table 1). IND, IAA, I-Ald, and IA were detected in all subjects, while ILA, TAM, SK and IPA were found in 95, 90, and 81% of subjects, respectively. Two metabolites, indoxyl-3-sulfate (I3S), and indole acetate (IAc), were not detected in any fecal sample. Wide variation was observed for all measured TMs both individually and as a collective (Figure 3A; Table S4). IND had the highest relative concentration across all samples, followed by IAA, SK, and I-Ald, which together constituted 95% of the total TMs measured (Figure 3B). While IND was the most abundant TM overall, a finding consistent with previous reports for human fecal samples (15, reviewed in 16), another TM dominated samples from eight subjects: SK (n=4 subjects), TAM (n=3), and I-Ald (n=1). In one additional subject, IND was co-dominant with IAA. These variations in total and individual TM levels among healthy adults suggest a redundancy of function among TMs or with other microbial metabolites. Total TM concentrations (Figure 3A) showed that 71.4% had total TM values in the 20-60 nmol/g range (Figure 3C).

**Figure 3.**
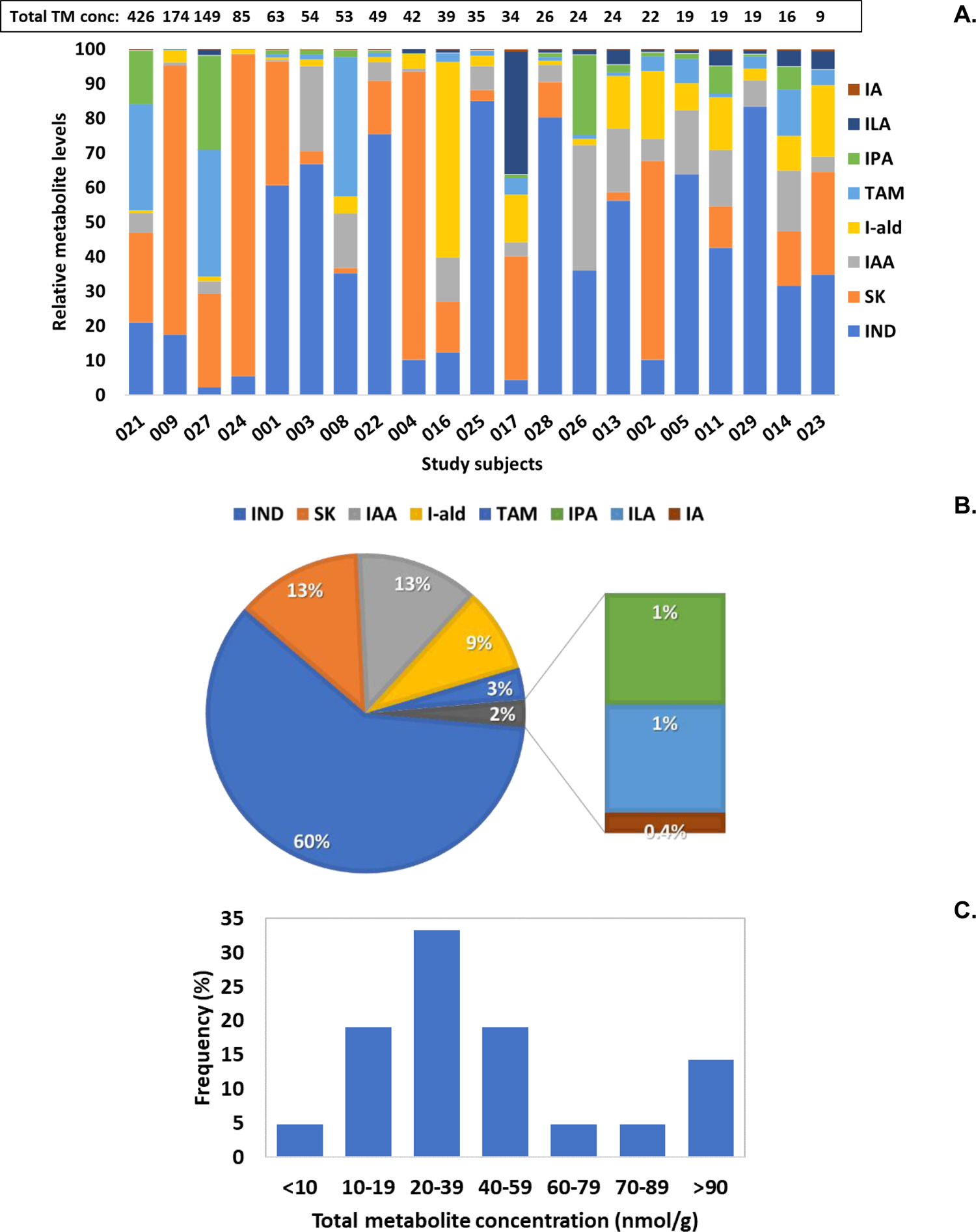
Tryptophan metabolites (TM, n=8) in stool samples from 21 healthy adults. A. TM levels are depicted as relative levels (%) per subject. Total TM levels were summed for each subject and are indicated above the figure. B. Relative levels of metabolite (median nmol/g). C. Frequency distribution of total metabolite concentrations (nmol/g). Abbreviations for TMs can be found in. Table 1.**Figure 4**.

**Figure 4.**
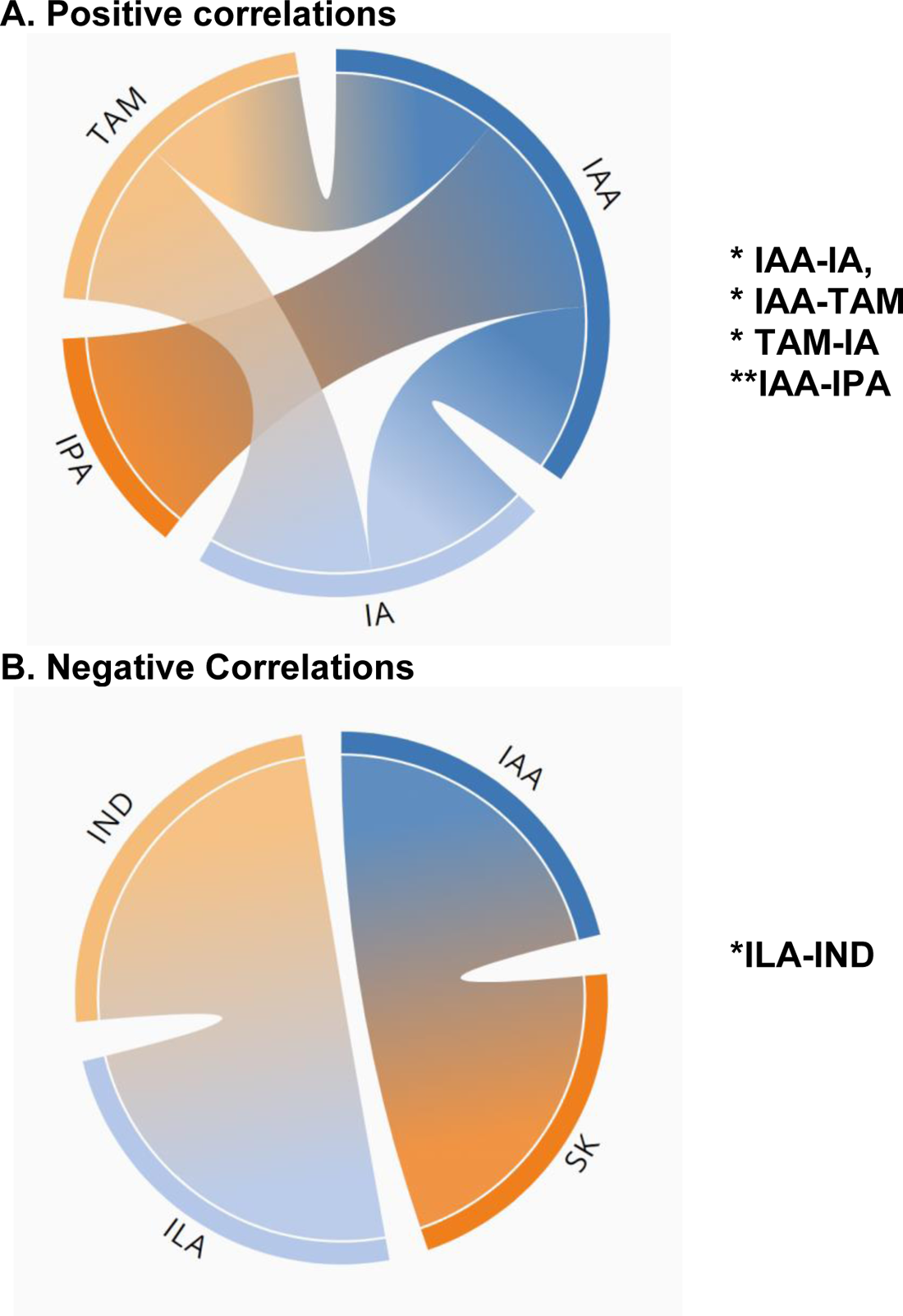
Significant relationships among tryptophan metabolites (nmol/g) in fecal samples from 21 healthy adults. Positive (A) and negative (B) correlations are shown. Data were generated using Spearman rank correlation (r); statistical significance (*p=<0.05 or **p= <0.01) is indicated.

Potential associations between TM levels and sex, race/ethnicity, or age were examined. TM levels between the sexes showed a significant difference (p=0.021, Mann-Whitney statistic) for ILA only. Males (n=8) had median ILA levels 8.5-fold higher (3.16 vs 0.37 nmol/g) than females (n=13). However, this difference depended on including data from non-White individuals (three Hispanics and one Black). When the comparison was limited to White participants (female, n=5; male, n=7), no significant differences were found for any TM alone or total TM levels. TM levels were then compared among the three race/ethnic groups (Black, n=6; Hispanic, n=3; White, n=12), and no significant associations were found. Further examination included females only (Black, n=6; White, n=5) given the limited numbers of Hispanics (n=3) and Black males (n=0). In that analysis, Black females had a 3.5-fold increase (Mann Whitney statistic, p= 0.017) in I-Ald levels, while none of the other TMs were significantly different. Lastly, the potential effect age was evaluated for the three race/ethnicity groups. Overall, no significant associations for TM levels were found. However, when sex was taken into consideration, IND in males (but not females) increased significantly (r^2^= 0.972) with age (range, 24-35 years).

Given the interconnectedness of metabolites in the indole pathway, we performed a correlation analysis among TM levels, which revealed several significant relationships (Figure 5, Table S5). Significant associations were: IAA with IPA (p= 0.002), IA (p= 0.019), and TAM (p= 0.021); TAM with IA (p= 0.014) (positive associations); and IND with ILA (p=0.048, negative association). Interestingly, SK and ILA, both generated solely through bacterial metabolism, were the only metabolites that did not significantly correlate with any other TM, although a negative association (p= 0.086) between IAA and SK was noted.

**Figure 5.**
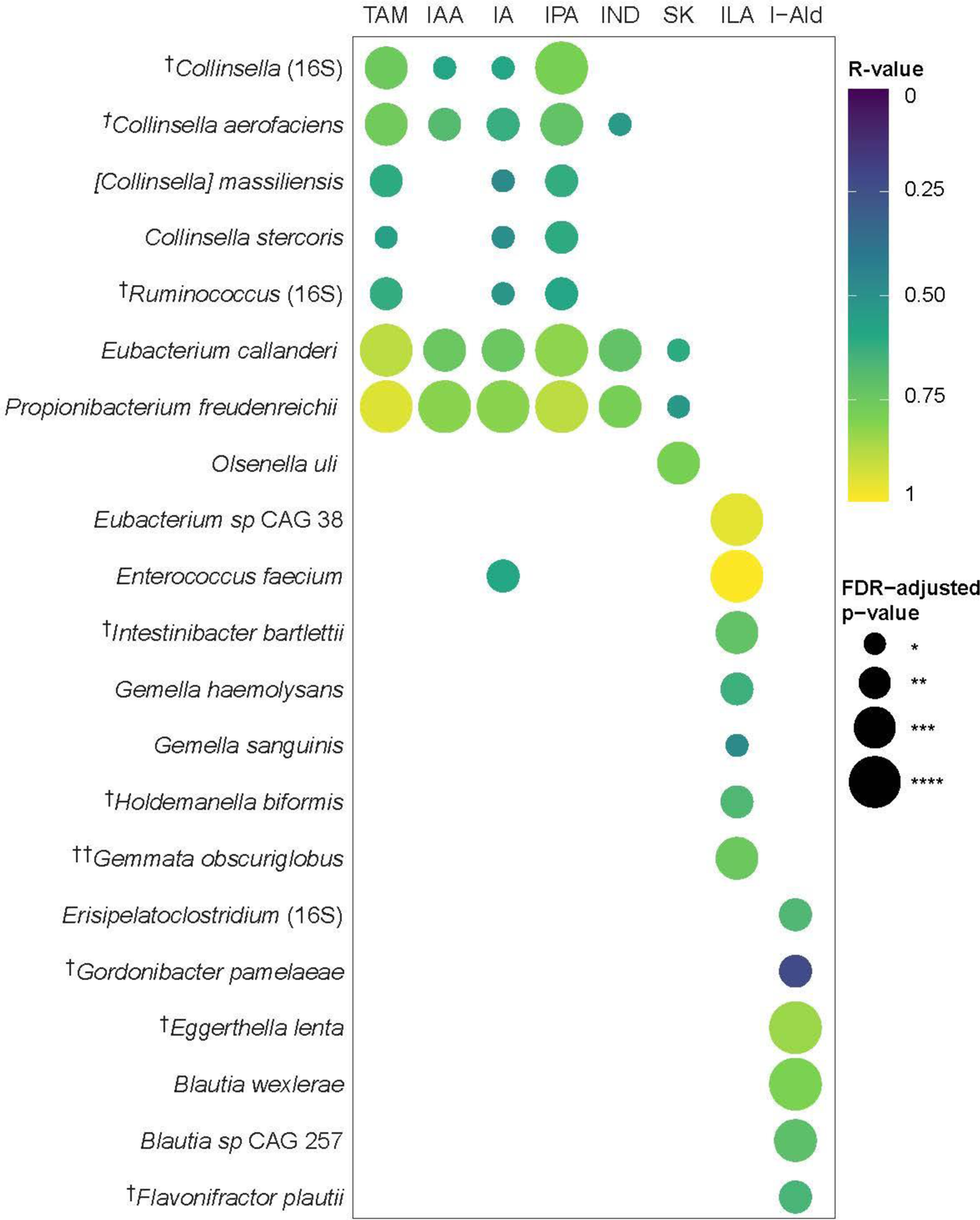
Linear regression correlations of bacterial taxa (% relative abundance) and tryptophan metabolite concentrations (nmol/g) in fecal samples from healthy adults. Taxa with a >20% occupancy and an FDR-adjusted p <0.05 are shown. Asterisks indicate taxa capable of producing IAA (†) or IND (††).

### Correlating microbiota and tryptophan metabolites

Due to the age of our stool samples, we first evaluated their bacterial content at the phylum level using 16S data for consistency with published reports. Microbial recovery from historic samples was evaluated at the phylum level. Ten (10) phyla were identified, the four most abundant accounting for >94% of total relative abundance. Compared to published reports in other healthy populations (25–31), we noted a substantial reduction in the relative abundance of Bacteroides in our samples. An apparent over-representation of Actinobacteria was also observed and may be due to compositionality effects related to the loss of Bacterioidetes, a phylum known to be more greatly impacted by frozen storage time and freeze-thaw cycles (30, 32). The relative abundance of other phyla did not vary significantly from expected levels (Table S3).

We surmised that TM concentrations (nmol/g) would vary with the abundance (%RA) of their microbial sources. Thus, we generated regression correlations (r) and FDR-adjusted p values for each metabolite and microbial species with >20% occupancy. In total, twenty-one (21) taxa were significantly correlated with one or more TMs (FDR-adjusted p <0.05; Figure 6), and each TM was associated with more than one taxon.

**Figure 6.**
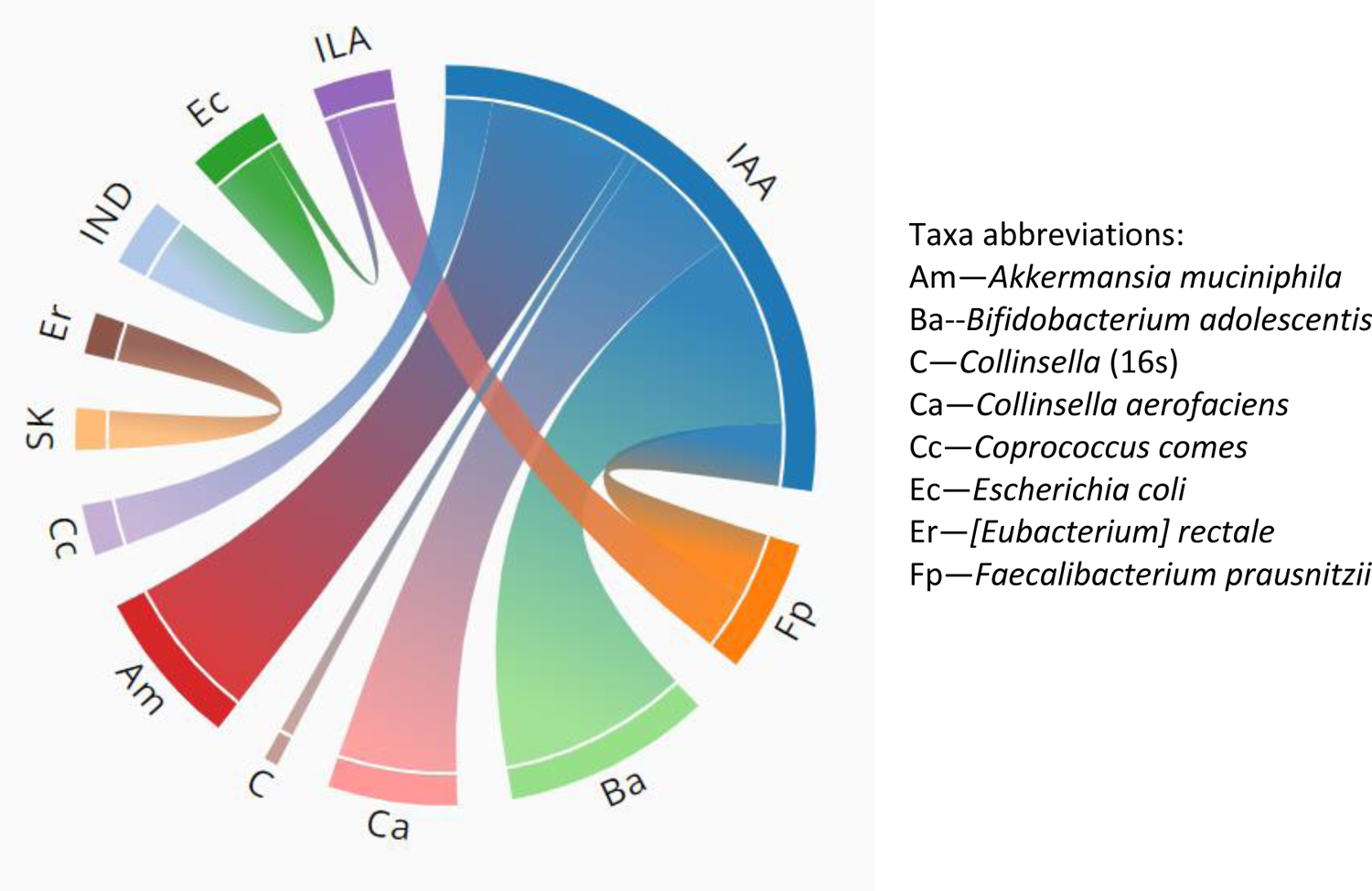
Predicted bacterial sources of tryptophan metabolites found in fecal samples from healthy adults. Mean prediction scores (see supplement 8) are shown for taxa with: >0.1 percent relative abundance (%RA); occupancy ≥5; and metabolite production probability)(MPP) >0.5.

Three distinct sets of bacterial taxa were evident: a set (n=7) dominated by *Collinsella* species, correlated with 3-6 TMs; and two taxa sets (n=7, n=6) each correlated with one TM, the exception being *Enterococcus faecium* which correlated with two. Interestingly, *Collinsella aerofaciens* was the only species that positively correlated with a metabolite (IAA) and had an MPP >0.5. A heat map illustrates the relative abundance of significantly correlated taxa (Table 2, Table S6) and is shaded for %RA >1. Of the fourteen (14) taxa in this category, six (6) taxa had occupancies of 5-19. *Ruminococcus*, *Blautia wexleri,* and *Collinsella aerofaciens* were in highest abundance (>10%) in multiple samples, suggesting a key role in tryptophan metabolism. It should be noted, however, that even low abundance taxa may be important since samples contain multiple TM-producing taxa that may act in concert.

**Table 2.**
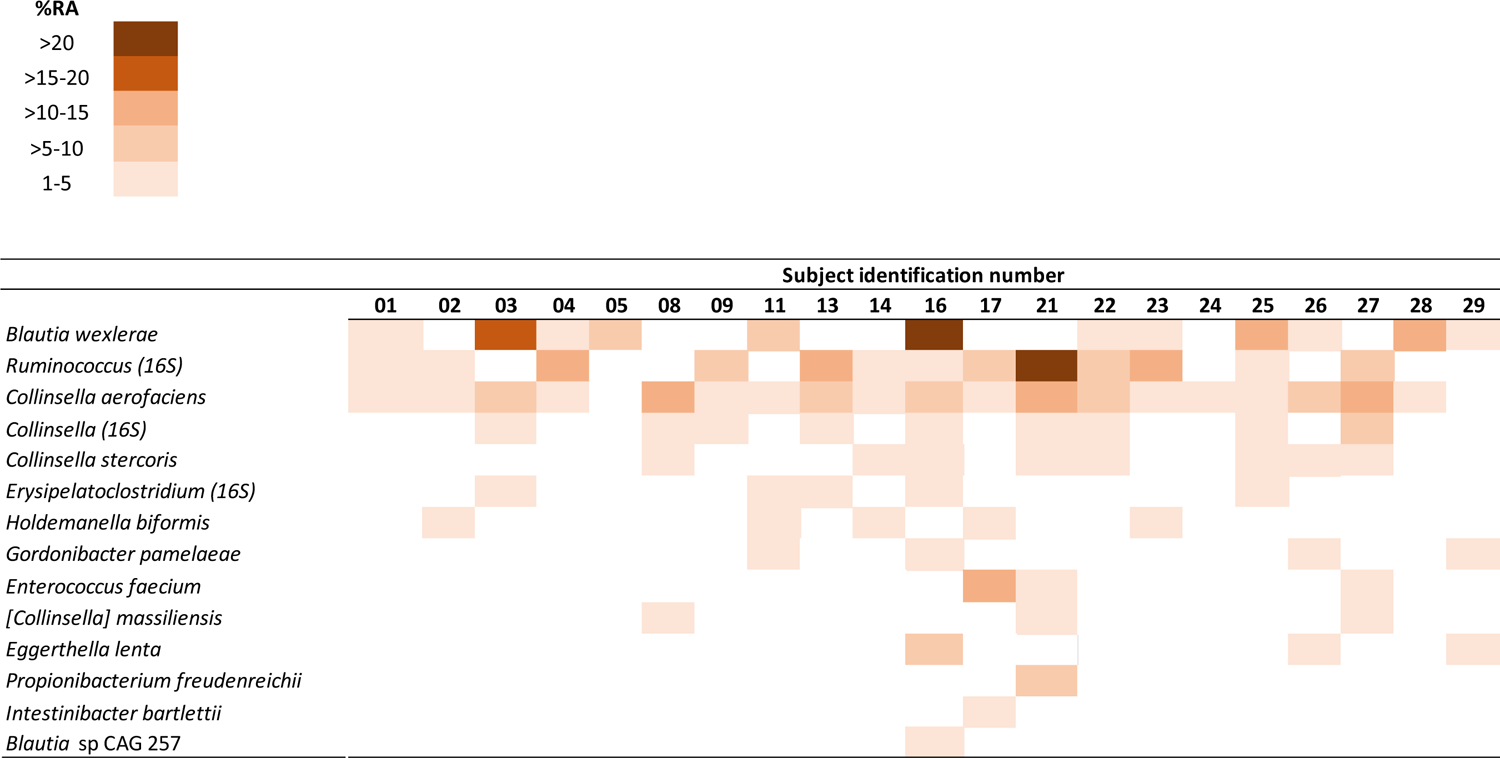
Heat map of percent relative abundance (%RA) for taxa that are significantly correlated with one or more tryptophan metabolites. Color scale represents relative abundance (%RA ≥1) for each subject.

### Predicting microbial sources of metabolites

A recent interactive website (www.TrpNet.ca) assigns probabilities to all known TM producers and allows for predictions based on taxon abundance in any biological sample. Twenty-four (24) species with metabolite-producing probabilities (MPPs) of >0.5 were identified in our study samples. Using this criterion, predictions for each sample were made for seven TMs (Figure 6, Table S7), resulting in prediction scores (MPP X %RA) ranging from 1-32. The highest number of taxa (n=20) were predicted for IAA production, and the most likely contributors by prediction score and occupancy were *Collinsella aerofaciens*, *Bifidobacterium adolescentis*, *Akkermansia muciniphila*, *Fecalibacterium prausnitzii, and Coprococcus comes*. Further, each fecal sample contained at least three taxa that could contribute to the observed IAA concentrations. Predictions for other TMs included five taxa for IND, two for ILA, and one each for SK and I-Ald, and all had <30% occupancy. No microbial source was found for TAM (data not shown), thus most, if not all, the observed TAM concentrations in the samples may have originated from the host. We then went one step further in the analysis by comparing prediction scores to observed TM levels for each of the taxon and TM combinations (Figure 7). Strong trends were seen with *Collinsella aerofaciens* versus IAA (r^2^= 0.6586) or total TM levels (r^2^= 0.8915). No correlation for any other taxon and TM combination exceeded an r^2^ of 0.33.

**Figure 7.**
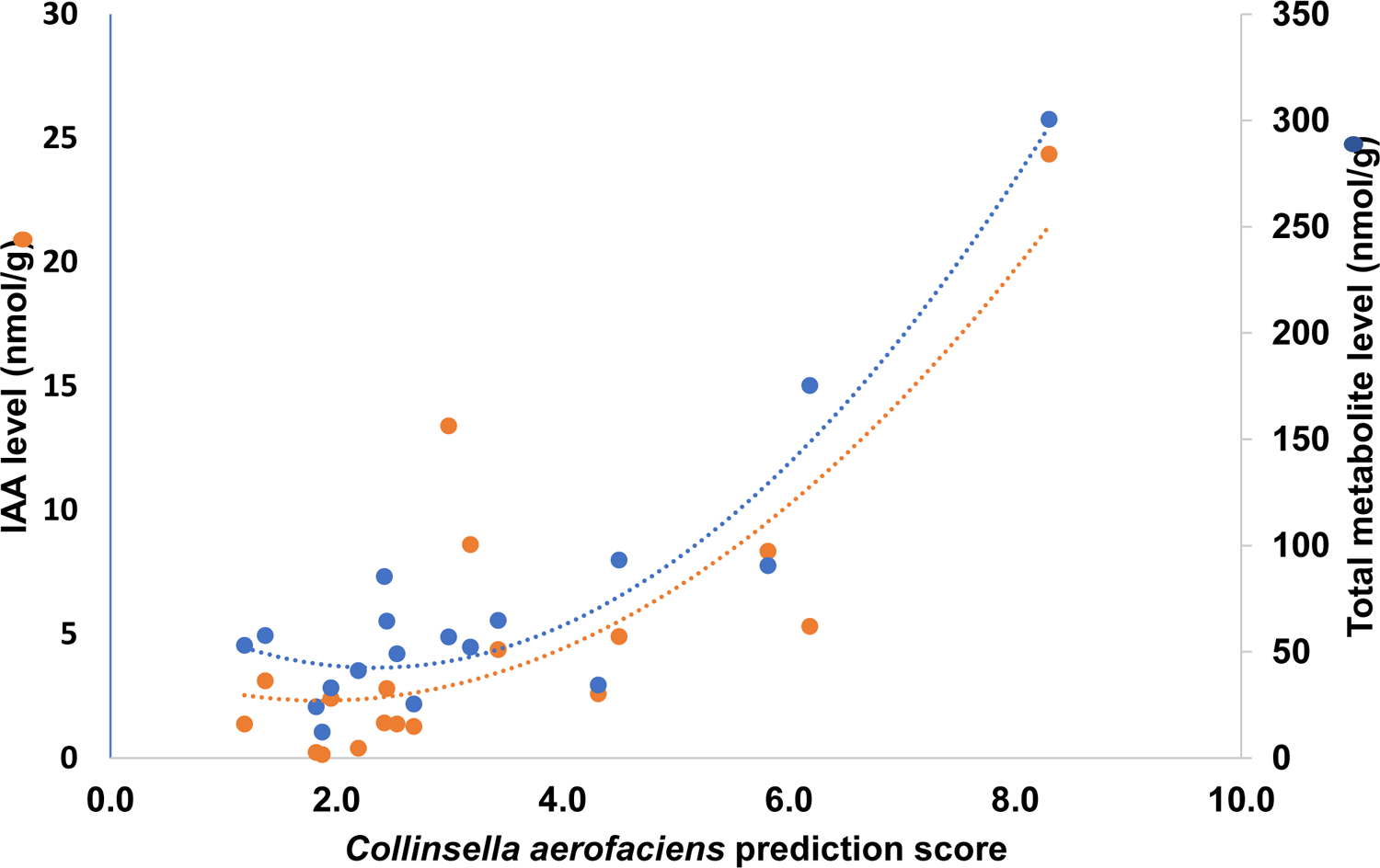
Correlation of fecal tryptophan metabolite concentrations with metabolite prediction score. Each marker represents individual data from a total of 21 healthy adults. Polynomial (2^nd^ order) curves of best fit are shown (dotted lines) for IAA levels (orange, r^2^ = 0.6586) and total levels of 8 tryptophan metabolites (blue, r^2^ = 0.8915).

### Summary of findings

Taken together, the data support the notion that multiple microbial species form a network of interactions involved in producing and maintaining TM levels needed for gut homeostasis. However, IAA and *Collinsella aerofaciens*, stand out as important components of the pathway. IAA was noteworthy in that it was: in high overall concentration; significantly correlated with three other TMs (TAM, IA, IPA); occupied a key position in the pathway; and had the largest number of taxa identified as predicted IAA sources. Among these taxa, *Collinsella aerofaciens* was the only one with all of the following characteristics: 100% occupancy; high MPP (0.516) for IAA production; high relative abundance (mean RA 5.5%); predicted IAA source in 85.7% of subjects; and positive trends between PS and IAA or total TM levels. In a “proof of concept” exercise, we constructed a hypothetical scenario in which *Collinsella aerofaciens* and IAA might “drive” the pathway toward metabolites that are important in gut homeostasis (Figure 8). TAM and/or IND arising from the host and/or microbial sources, respectively “feed” substrates to *Collinsella aerofaciens* (and the other indicated sources) to produce IAA. Increased IAA levels (which were positively associated with IA and IPA) could increase IA and IPA concentrations to achieve biologically active levels. Previous studies have shown that IA and IPA help to maintain the epithelial barrier, act as anti-inflammatory signals, and are strong antioxidants. In summary, this scenario illustrates an approach that combines TM measurements and sequencing data from fecal samples with metabolite production probabilities in a way that links tryptophan metabolites to particular taxa and further predicts likely microbial metabolite sources.

**Figure 8.**
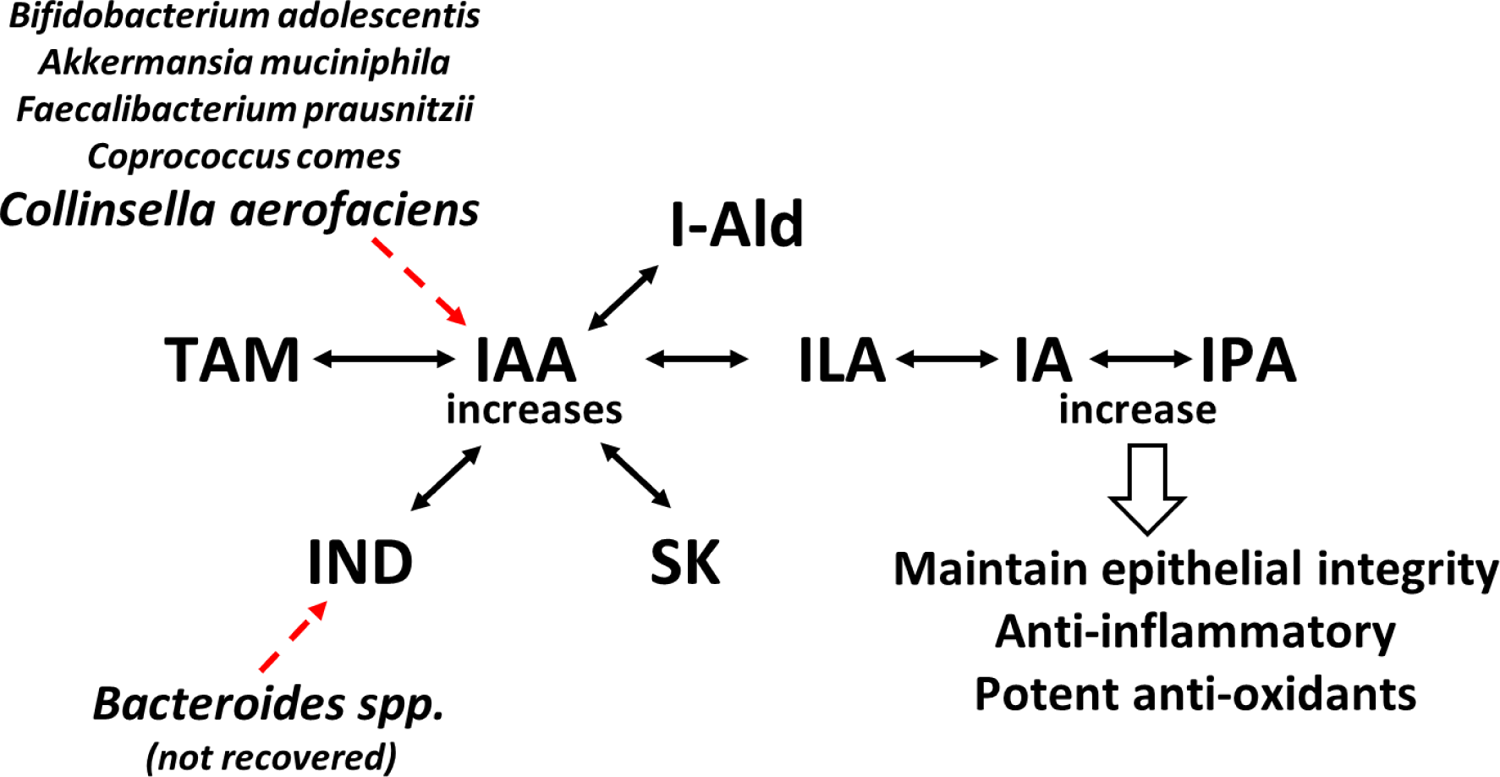
Major features of a hypothetical scenario leading to important functional outcomes for the host. The pathway is based on analyses of TM interactions, correlations between relative levels of taxa and TMs, and predictions of potential bacterial sources.

## DISCUSSION

The intestinal microbial community is shaped by diet, environment, age, ethnicity, and the host immune response, among other factors. In a “healthy” microbiome, necessary functions can be fulfilled by multiple species, thus allowing adaptation to new environments or conditions. This notion extends to molecular signals, including TMs, produced and secreted to maintain homeostasis. Thus, the wide variations that we found in TM levels among healthy individuals were not unexpected or unprecedented. Our study focused on eight TMs of the indole pathway and, therefore, represent only a portion, albeit an important one, of the greater metabolic “whole”. We fully recognize the complexity of interactions regulating myriad gut functions, which are known to involve both metabolites of microbial and host pathways, many presumably acting in concert.

Intestinal microbiota are the primary, and often sole, sources of fecal TMs. Many TM-producing taxa are known, but heretofore the tools for linking fecal TM levels to specific taxa were not readily available for studying complex biological samples. We were able to capture the most complete picture of the microbiota present in our samples by using 16S and WGS data, which rely on different sequencing techniques and databases to assign taxonomic identities. In general, 16S yields a broader array of bacterial taxa but lacks species specificity, whereas WGS provides specificity but is less comprehensive.

### TM measurements and relationships

We included ten fecal metabolites with previously recognized intestinal functions contributing to gut homeostasis as well as having anti-inflammatory roles in extraintestinal tissues. Eight of these metabolites were found in 81 – 100% of samples from healthy subjects but varied widely from person to person (Figure 3A). Many factors likely contribute to the observed variations, but diet is recognized as playing a major role in shaping the gut microbial community (31; reviewed in 33-34). No dietary information was collected on individuals in the study. IND, SK, and IAA were in highest overall concentrations (Figure 3B). All TM levels we measured add significantly to the limited literature, but to our knowledge, the ILA, IPA, and IA concentrations represent the first reports from human fecal samples. Published human fecal TM levels vary with method and population. In comparison, our study was generally consistent with a recent report using the same methodology (15). Not surprisingly, we did not detect I3S or IAc in any samples. I3S, produced in the liver, is associated with pathological conditions such as chronic kidney disease (reviewed in 35), and indole acetate (IAc) is the transient base conjugate of IAA hydrolyzed by host and bacterial esterases. We also note that MS-based methods typically yielded lower TM concentrations than photometric methods. The reasons for this difference were unclear but may be related to method specificity or recovery efficiencies of sample preparation steps.

As expected, several significant associations were found among the TMs (Figure 4), but IAA appeared to have a key role in the pathway since it was significantly associated with three other metabolites (TAM, IA, and IPA). Further, IAA is a required step for two other significantly associated metabolite pairs (IND and ILA, TAM and IA). Exploration of relationships between TM levels with demographic categories was limited by sample size and representation. No significant association was found between White males and females. However, when all samples were included, males (87.5% White) had significantly higher ILA levels (p=0.004). Race and age were also evaluated and showed a difference in I-Ald (higher in Black versus White females, p=0.017) and IND increases with age (males, r^2^=0.972). However, these results are preliminary and require verification in larger populations.

Armed with TM quantitation, our attention turned to identifying their potential sources. To that end, we assumed that higher TM levels would correspond with higher relative abundance, and correlation analyses did indeed point to multiple taxa associated with each TM (Figure 5). However, significant associations were almost always between a TM and a taxon not known to produce it. Of nine IAA- or IND-producing taxa, only *Collinsella* (16S) and *Collinsella aerofaciens* correlated with their product, IAA. Interestingly, another study that showed IAA could alter concentrations of many microbe-specific metabolites (36). Further, twelve non-TM-producing taxa were also significantly correlated with ≥1 TMs, suggesting that their relative abundances might influence, or be influenced by, TM production, perhaps via a metabolic product not included in the study.

Until recently, microbial metabolite production was established by assessment of supernatants from purified cultures and/or *in silico* analysis of pathway genes (7, 21, 22). As a result, bacterial species were typically dichotomized as metabolite producers or non-producers. More recently, information gathered from all known public sources has been incorporated into a Bayesian logistic regression model, which assigns a metabolite production probability (MPP) for each step of the tryptophan pathway (23). With this information, we identified bacterial species found in our dataset that were likely sources of the observed fecal TM levels (Figure 7). While several potential sources were found for IAA, that was not the case for other metabolites. It is possible that cutoffs used in the analyses (i.e. MPP >0.5, %RA >1) excluded low abundance taxa that each contributed minimally but together might have accounted for the observed TM levels. Lastly, the lack of higher prediction scores for IND and SK sources was not unexpected due to the low recovery of *Bacteroides* spp., a rich source for these metabolites. Of further note, while prediction scores can provide insight into likely TM sources, they do not necessarily predict the quantity of TMs produced. Indeed, the amount of metabolite produced by a particular taxon depends on such factors as having the necessary metabolic machinery, abundance, substrate availability, and regulatory signals from the microbial community and/or the host, among others.

### Limitations

There are two main limitations in our study: the recovery of microbial taxa and metabolites in historic fecal samples and the number of samples included in the study. Long-term cryopreservation of bacteria is common in biobanking, and much has been learned over the past two decades about enhancing the recovery of bacteria from stored samples (reviewed in 37). Fecal samples used in the study were stored at −80C years before post-storage recovery techniques had been studied to any great degree. Further, when the samples were collected, sequencing techniques were not available to establish baselines. Microbial recovery was comparable to other studies in that Firmicutes, Actinobacteria, Verrucomicrobia, and Proteobacteria were well preserved (reviewed in 37), but Bacteroidetes were greatly decreased. The relative loss of Bacteroidetes species was unfortunate since it is a major source of IND and SK and compromised our ability to confirm the role of *Bacteroides spp.* as a significant source of these compounds. On the other hand, by limiting *Bacteroides* abundance, compositionality effects may have enhanced our ability to identify important relationships with lower abundance taxa. Even with these limitations, our findings demonstrate that long-term cryopreservation of fecal samples can be used to provide useful information. Lastly, the number of fecal samples and TMs studied was limited by the expense involved in mass spectroscopy but nevertheless exceeds the sample sizes in many publications and significantly adds to the existing literature.

Finally, we then used our collective observations in a hypothetical scenario suggesting that *Collinsella aerofaciens* (and four other taxa) play a leading role in producing high IAA levels, driving the pathway toward increased IA and IPA, which then contribute to maintaining epithelial integrity, acting as reactive oxygen scavengers, and decreasing the gut inflammatory response.

In summary, using an approach such as the one we have presented here can be useful in selecting microbial species for testing in animal models, thus supporting a rational approach to developing “next-generation” probiotics. Efforts using selected combinations of microbial species designed to address specific dysbiosis are in the early stages (38, 39) but promise a new era in treating a variety of serious health conditions affecting a large portion of the population.

## ACKNOWLEDGEMENTS

This work was supported by The National Institutes of Health, U19AI144297,(Joseph Petrosino, program PI), and Pablo C. Okhuysen and Cynthia L Chappell, (subcontract co-PIs). We wish to thank Srikar Ranga and Gabriella Rodriguez for their technical assistance.

## CO-AUTHOR ROLES

Cynthia Chappell – conceptualization, investigation, formal analysis, project administration, resources, validation, visualization, writing Kristi Hoffman -- formal analysis, writing Philip L. Lorenzi -- methodology, resources, validation Lin Tan. Investigation, methodology Joseph Petrosino – funding acquisition, project administration, resources Richard A. Gibbs– funding acquisition, project administration, resources Donna M. Muzny – data curation, formal analysis Harsha Doddapaneni – data curation, formal analysis Matthew C. Ross – data curation, formal analysis Vipin K. Menon – data curation, formal analysis Anil Surathur -- data curation, formal analysis Sara J. Javornik Cregeen -- formal analysis, visualization Anaid G. Reyes – validation Pablo Okhuysen—conceptualization, project administration, resources, writing

## DISCLAIMERS

The funders had no role in study design, data collection and interpretation, or the decision to submit the work for publication. None of the authors have a financial or other relationship with TrpNet.

## Notes

### Competing Interest Statement

The authors have declared no competing interest.

### Summary of Updates

No revision has been done. The paper was transferred from one ASM journal to another.

