## Supplemental Data for "Tryptophan Metabolites And Their Predicted Microbial Sources In Fecal Samples Of Healthy Individuals"

**Chappell et al.**

### Supplementary Data

***Table S1. Published studies measuring one or more tryptophan metabolites in stool samples. All publications except Dong et al., 2020 were included in a review by Anderson et al., 2021 from which the following table is adapted. All metabolite measurements have been converted to ug/g.***

| Study | Method | IND | SK | IAA | TAM | I-ald | IA | IPA |
|---|---|---|---|---|---|---|---|---|
| **Human** | | | | | | | | |
| Bergeim, 1917 | Color | 26.0 | | | | | | |
| Hibbard et al., 2019 | Color | 617.0 | | | | | | |
| Karlin et al., 1985 | GC | 58.0 | 34.5 | | | | | |
| Zuccato et al., 1993 | GC | 101.0 | 33.9 | | | | | |
| Darkoh et al., 2015 | Photo | 304.0 | | | | | | |
| Chappell et al., 2016 | Photo | 340.0 | | | | | | |
| Zhao et al., 2018 | Photo | 35.0 | | | | | | |
| Lamas et al., 2016 | LC/MS | | | 0.894 | | | | |
| Shi et al., 2020 | LC/FD | 12.0 | 1.0 | | | | | |
| Dehnhard et al., 1991 | LC/UV | | 15.5 | | | | | |
| DeFilipis et al., 2016 | GC/MS | 11.4 | | | | | | |
| Dong et al., 2020 | GC/MS | 0.057 | 0.001 | 0.004 | 0.001 | 9.0E-05 | <LOD | |
| **Mouse** | | | | | | | | |
| Jin et al., 2014 | LC/MS | 42 | | 2.6 | 1.6 | | | |
| Zheng et al., 2020 | LC/MS | | | 1100.0 | | | | 490 |
| Dong et al., 2020 | GC/MS | 0.001 | | 0.001 | 8.0E-04 | 7.0E-05 | 1.10E-05 | |
| **Rat** | | | | | | | | |
| Anderson et al., 1975 | TLC | 6.4 | 0.2 | 2.1 | 0.67 | | | 2.5 |
| Lamas et al., 2016 | LC/MS | | | 0.175 | | | | |
| Zeng et al., 2019 | LC/MS | | | 0.935 | | | | 0.302 |
| Liu et al., 2018 | LC/MS | 1.7 | | | | | | |

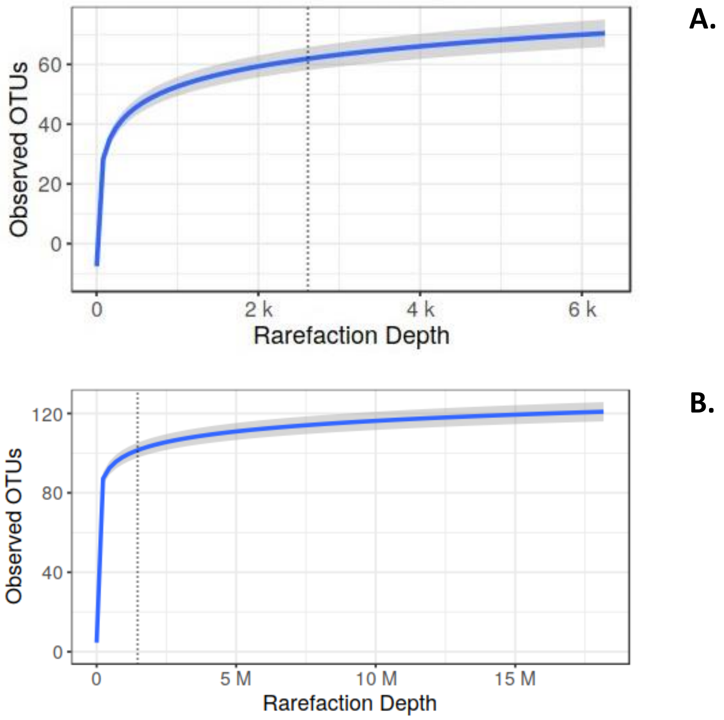


***Figure S1. 16S (A) and whole genome sequencing (B) rarefaction curves for observed OTUs. All samples were retained at rarefaction depths of 2272 (A) and 1451921 (B).***

***Table S2. Demographic information for the study population.***

| Subject ID | Age | Sex | Ethnicity |
|---|---|---|---|
| **001** | 35 | Male | White |
| **002** | 44 | Female | Black |
| **003** | 40 | Female | Black |
| **004** | 24 | Male | White |
| **005** | 41 | Female | Black |
| **008** | 37 | Female | Black |
| **009** | 42 | Female | Black |
| **011** | 29 | Male | White |
| **013** | 30 | Male | White |
| **014** | 28 | Male | White |
| **016** | 24 | Male | White |
| **017** | 27 | Male | Hispanic |
| **021** | 45 | Female | Hispanic |
| **022** | 25 | Female | White |
| **023** | 22 | Female | Black |
| **024** | 37 | Female | White |
| **025** | 25 | Female | White |
| **026** | 24 | Female | White |
| **027** | 29 | Male | White |
| **028** | 37 | Female | Hispanic |
| **029** | 29 | Female | White |

***Table S3. Relative abundance of phyla from published studies of healthy individuals. Data were calculated from 16S sequencing of fecal samples.***

| | Hseih et al., 2016 | Eckberg, 2005 | Zhu et al., 2020 | Sartor, 2008 | Frank et al., 2007* | Arumugam et al., 2011 | Yatsunenko et al., 2012 | Median (7 studies) |
|---|---|---|---|---|---|---|---|---|
| **Firmicutes** | 61.2-85.7 | 76.2 | 73.3-76.0 | 64 | 55 | 36 | 8.45 | 64.0 |
| **Actinobacteria** | 0.5-16.3 | 1.8 | 7.8-8.3 | 4 | 5 | 5 | 4.15 | 5.0 |
| **Bacteroidetes** | 7.1-29.6 | 16.5 | 14.2-16.7 | 23 | 28 | 23 | 83.7 | 23 |
| **Proteobacteria** | 0.2-2.0 | 1.3 | 1.2 | 8 | 12 | <1 | -- | 1.3 |
| **Verrucomicrobia** | -- | 0.2 | 0.54-0.62 | -- | -- | <1 | -- | 0.2 |
| **Other** | -- | | 0.18 | 1 | | | 3.75 | 1.6 |

*Exact data not available; estimated from published figure

***Table S4. Descriptive statistics for eight tryptophan metabolites quantitated from fecal samples of 21 healthy adults. Means, standard deviations, and medians are expressed as nmol/g stool.***

| | Tryptophan metabolites | | | | | | | |
|---|---|---|---|---|---|---|---|---|
| | **IND** | **SK** | **IAA** | **I-Ald** | **TAM** | **IPA** | **ILA** | **IA** |
| **# of Subjects:** | 21 | 19 | 21 | 21 | 19 | 17 | 20 | 21 |
| Mean | 18.4 | 22.7 | 4.4 | 3.0 | 10.4 | 5.7 | 1.1 | 0.1 |
| SD | 20.5 | 38.7 | 5.7 | 4.6 | 30.3 | 16.4 | 2.6 | 0.1 |
| Median | 12.4 | 2.7 | 2.6 | 1.8 | 0.7 | 0.3 | 0.3 | 0.1 |

***Table S5. Spearman rank correlation (r) matrix for tryptophan metabolites (nmol/g) in fecal samples of 21 healthy adults. Significant values (p= <0.05, red) and trends (p >0.05 - <0.1, blue) are indicated.***

| | | Tryptophan metabolite | | | | | | |
|---|---|---|---|---|---|---|---|---|
| | | **IA** | **IPA** | **SK** | **ILA** | **IND** | **I-Ald** | **TAM** |
| **IAA** | r = | 0.5075 | 0.6352 | -0.3836 | 0.1812 | 0.2429 | 0.174 | 0.5015 |
| | p = | **0.0189** | **0.002** | **0.086** | 0.4318 | 0.2888 | 0.4506 | **0.0206** |
| **IA** | r = | | 0.412 | -0.1067 | 0.0871 | 0.2313 | -0.1072 | 0.5271 |
| | p = | | **0.0635** | 0.6453 | 0.7074 | 0.313 | 0.6437 | **0.0141** |
| **IPA** | r = | | | -0.107 | 0.3617 | 0.258 | -0.0736 | 0.3731 |
| | p = | | | 0.6444 | 0.1072 | 0.2589 | 0.7511 | **0.0958** |
| **SK** | r = | | | | 0.1405 | -0.0819 | 0.2926 | 0.0416 |
| | p = | | | | 0.5436 | 0.7241 | 0.1981 | 0.8578 |
| **ILA** | r = | | | | | -0.4359 | 0.4144 | 0.1787 |
| | p = | | | | | **0.0483** | **0.0618** | 0.4384 |
| **IND** | r = | | | | | | -0.3805 | 0.0474 |
| | p = | | | | | | **0.0888** | 0.8383 |
| **I-Ald** | r = | | | | | | | 0.2787 |
| | p = | | | | | | | 0.2213 |

***Table S6. Heat map of taxa that are significantly correlated with one or more tryptophan metabolites. Values represent relative abundance (RA >0.1%) for each subject; color scale is indicated for RA ≥1%.***

| %RA |
|---|
| >20 |
| >15-20 |
| >10-15 |
| >5-10 |
| 1-5 |

| | Subject identification number | | | | | | | | | | | | | | | | | | | | |
|---|---|---|---|---|---|---|---|---|---|---|---|---|---|---|---|---|---|---|---|---|---|
| | **01** | **02** | **03** | **04** | **05** | **08** | **09** | **11** | **13** | **14** | **16** | **17** | **21** | **22** | **23** | **24** | **25** | **26** | **27** | **28** | **29** |
| *Blautia wexlerae* | 2.3 | 0.5 | 17.4 | 1.3 | 10.3 | 0.8 | 0.2 | 8.3 | 0.4 | 0.8 | 50.4 | 0.4 | 0.1 | 1.0 | 1.1 | 0.1 | 11.5 | 2.8 | | 11.4 | 1.1 |
| *Ruminococcus (16S)* | 3.7 | 1.1 | | 13 | | 0.6 | 7.8 | | 12 | 1.5 | 4.53 | 6.6 | 23.2 | 6.6 | 14.3 | 0.8 | 2.1 | 0.4 | 5.5 | 0.3 | |
| *Collinsella aerofaciens* | 3.5 | 2.3 | 5.8 | 0.1 | | 11.3 | 4.9 | 2.7 | 6.6 | 4.7 | 8.7 | 4.7 | 16.1 | 8.4 | 4.2 | 3.6 | 3.8 | 6.2 | 12.0 | 5.2 | |
| *Collinsella (16S)* | 0.2 | 0.7 | 1.1 | | | 1.5 | 1.5 | 0.3 | 1.7 | 0.2 | 3.5 | 0.4 | 5.0 | 2.8 | 0.8 | 0.5 | 1.1 | 0.3 | 5.1 | 0.6 | 0.0 |
| *Collinsella stercoris* | 0.6 | 0.6 | 0.7 | | | 1.0 | 0.6 | 0.3 | 0.8 | 1.0 | 1.1 | 0.7 | 1.5 | 1.1 | 0.6 | 0.5 | 1.3 | 3.2 | 2.1 | 0.7 | |
| *Erysipelatoclostridium (16S)* | 0.9 | | 1.9 | 0.1 | 0.4 | | 0.2 | 1.8 | 1.5 | | 3.5 | | | | | 0.3 | 1.2 | 0.2 | | | 0.2 |
| *Holdemanella biformis* | | 2.4 | | | | 0.3 | | 1.8 | 0.3 | 2.5 | | 3.8 | | | 1.0 | | | | | | |
| *Gordonibacter pamelaeae* | 0.2 | 0.3 | 0.6 | 0.1 | | 0.2 | 0.1 | 1.3 | 0.1 | 0.4 | 4.0 | 0.3 | 0.9 | 0.4 | 0.1 | 0.1 | 0.9 | 1.8 | | | 2.2 |
| *Enterococcus faecium* | | | | | | | | | | | | 15.0 | 2.0 | | | | | | 1.2 | | |
| *[Collinsella] massiliensis* | 0.4 | 0.3 | 0.4 | 0.1 | | 1.0 | 0.4 | 0.3 | 0.5 | 0.6 | 0.9 | 0.5 | 1.0 | 0.6 | 0.4 | 0.3 | 0.4 | 0.9 | 1.3 | 0.4 | 0.1 |
| *Eggerthella lenta* | 0.1 | | 0.7 | 0.1 | | 0.1 | | 0.3 | | | 7.3 | | 0.7 | 0.4 | 0.1 | | 0.7 | 1.3 | | | 2.7 |
| *Propionibacterium freudenreichii* | | | | | | | | | | | | 0.1 | 5.1 | | | | | | | | |
| *Intestinibacter bartlettii* | 0.4 | | | 0.1 | 0.4 | 0.7 | | | 0.3 | 0.4 | | 1.7 | 0.2 | | | 0.1 | 0.4 | 0.4 | | | 0.1 |
| *Blautia* sp CAG 257 | | | 0.4 | | | | | | | | 1.3 | | | 0.1 | | | 0.1 | | | | 0.9 |

***Table S7. Predicted bacterial sources for seven tryptophan metabolites found in fecal samples from healthy subjects. All bacterial species with a metabolite production probability (MPP, www.TrpNet.ca) of >0.1 were included. Prediction scores (PS) for each subject and bacterial species were categorized (scale shown below) for likelihood of species as a metabolite source.***

| PS | | |
|---|---|---|
| >15 | | *Highly likely* |
| <15-7 | | |
| <7-3 | | |
| <3-1 | | |
| <1 | | *Unlikely* |

| | | Subject ID | | | | | | | | | | | | | | | | | | | | |
|---|---|---|---|---|---|---|---|---|---|---|---|---|---|---|---|---|---|---|---|---|---|---|
| **Bacterial species** | **MPP** | **01** | **02** | **03** | **04** | **05** | **08** | **09** | **11** | **13** | **14** | **16** | **17** | **21** | **22** | **23** | **24** | **25** | **26** | **27** | **28** | **29** |
| | | | | | | | | | | **Indole acetic acid (IAA)** | | | | | | | | | | | | |
| *Collinsella aerofaciens* | 0.516 | 1.8 | 1.2 | 3.0 | | | 5.8 | 2.5 | 1.4 | 3.4 | 2.4 | 4.5 | 2.4 | 8.3 | 4.3 | 2.2 | 1.9 | 1.9 | 3.2 | 6.2 | 2.7 | |
| *Bifidobacterium adolescentis* | 0.696 | 8.5 | 1.6 | | 10.1 | | 4.3 | | | | 9.8 | | 16.9 | | 10.2 | 5.0 | | | | 27.7 | | |
| *Akkermansia muciniphila* | 0.516 | | 5.5 | | 0.2 | 7.8 | | | 0.1 | 1.3 | | 2.6 | | | 0.1 | 6.3 | 32.0 | 2.2 | 13.2 | | 0.1 | |
| *Faecalibacterium prausnitzii* | 0.563 | 0.6 | 18.2 | 0.2 | 1.5 | | 0.1 | 0.8 | 1.6 | 0.7 | 0.8 | 0.1 | 0.1 | | 1.6 | 2.9 | 0.4 | | 1.6 | | 0.5 | 0.3 |
| *Coprococcus comes* | 0.516 | 2.7 | 1.2 | 2.2 | | | 1.6 | 2.1 | 3.4 | 0.6 | 0.8 | 1.9 | 0.5 | | 1.3 | 0.5 | | 0.4 | 0.2 | | 1.2 | 2.9 |
| *Bifidobacterium longum* | 0.172 | 2.8 | 0.1 | 0.4 | 4.9 | | 1.9 | 1.0 | 2.9 | 8.2 | 1.2 | | 1.9 | 0.6 | 0.3 | 0.2 | 0.1 | 0.3 | 1.1 | 4.3 | | 2.6 |
| *Blautia obeum* | 0.255 | 1.1 | 0.1 | 0.8 | 0.1 | | 0.6 | 1.0 | 0.4 | 0.3 | 0.3 | | 0.5 | | 1.4 | 0.1 | 0.1 | 8.1 | 2.3 | 0.1 | 3.5 | 0.3 |
| *[Ruminococcus] torques* | 0.245 | 0.7 | 0.9 | 1.4 | 0.2 | 1.7 | 0.7 | 2.7 | 0.2 | 0.6 | 3.5 | | 0.6 | 0.1 | 1.5 | 1.2 | 0.1 | 1.9 | | | 0.8 | |
| *Dorea longicatena* | 0.516 | 1.7 | 1.0 | 1.6 | 0.4 | | 1.9 | 0.8 | 2.5 | 0.5 | 0.1 | 0.2 | 1.7 | 0.0 | 0.8 | 2.1 | 0.5 | 1.7 | 1.6 | 0.2 | 1.8 | 1.3 |
| *Dorea formicigenerans* | 0.516 | 0.6 | 0.4 | 2.3 | | 2.2 | 1.4 | 0.7 | 0.4 | 0.1 | 0.7 | | 0.1 | 0.4 | 0.3 | 0.1 | 0.3 | 0.3 | 0.1 | 0.1 | 0.7 | 4.5 |
| *Ruminococcus lactaris* | 0.516 | | | 3.0 | | | | | 3.1 | 1.8 | 2.2 | | | | | | | | 0.1 | | 0.1 | |
| *Dorea* sp CAG 317 | 0.720 | | | 3.0 | | 6.9 | | 0.1 | | | | | | | | | | | | | | |
| *Escherichia coli* | 0.929 | 0.1 | | | 0.1 | | 0.1 | | | 0.2 | | | | | 5.6 | 0.1 | 3.8 | | | | | |
| *Coprococcus eutactus* | 0.516 | | 1.6 | | | | | | | | 0.8 | | 0.3 | | | 4.4 | | | | | 0.1 | |
| *Adlercreutzia equolifaciens* | 0.516 | 0.6 | 0.6 | 1.3 | 0.2 | | 0.2 | 0.1 | 1.0 | 0.2 | 0.5 | 0.3 | 0.2 | 0.2 | 0.3 | 0.1 | 0.2 | 0.8 | 0.5 | 0.3 | 0.4 | 0.1 |
| *[Eubacterium] rectale* | 0.245 | 2.4 | 0.4 | 1.5 | 0.2 | | | 0.1 | 1.6 | | 0.1 | 0.1 | | | | 1.0 | | | 0.3 | | | 0.2 |
| *Holdemanella biformis* | 0.516 | | 1.2 | | | | 0.1 | | 0.9 | 0.1 | 1.3 | | 2.0 | | | 0.5 | | | | 0.1 | 0.2 | |
| *Eubacterium* sp CAG 251 | 0.682 | 5.9 | | | | | | | | | 1.2 | | 0.5 | | 0.6 | | | | | 0.1 | | |
| *Eggerthella lenta* | 0.516 | 0.1 | | 0.4 | | | | | 0.2 | | | 3.8 | | 0.4 | 0.2 | | | 0.4 | 0.6 | | 0.2 | 1.4 |
| *Gordonibacter pamelaeae* | 0.516 | 0.1 | 0.2 | 0.3 | | | 0.1 | 0.1 | 0.7 | | 0.2 | 2.0 | 0.2 | 0.5 | 0.2 | 0.1 | | 0.5 | 0.9 | 0.3 | 0.2 | 1.1 |
| *Ruthenibacterium lactatiformans* | 0.516 | 0.1 | 0.1 | 1.0 | 0.1 | | | 0.1 | 0.2 | 0.3 | 0.1 | | | | 1.6 | 0.1 | 0.9 | | 0.5 | | | 0.4 |
| *Eubacterium ramulus* | 0.550 | | 0.3 | | | | | | 0.1 | | | | 0.0 | | 0.3 | | | 0.0 | 2.5 | | | |
| *Finegoldia magna* | 0.516 | | | | | | | | | | | | 3.1 | | | | | | | | | |
| *Ruminococcus bicirculans* | 0.516 | | 1.4 | | 0.5 | | | | | 0.1 | 0.5 | | | | | | | | | | | |
| *Bacteroides fragilis* | 0.836 | | 0.1 | 0.3 | | | | | | 0.8 | | | | | 1.1 | | | | 0.1 | | | |
| *Intestinibacter bartlettii* | 0.516 | 0.2 | | | | 0.2 | 0.4 | | | 0.1 | 0.2 | | 0.9 | 0.1 | | | 0.1 | 0.2 | 0.2 | | | 0.1 |
| *Olsenella scatoligenes* | 0.673 | 0.1 | 0.1 | | | | 0.1 | 0.1 | | 0.1 | 0.1 | 0.1 | 0.1 | 0.1 | 0.1 | 0.1 | 0.1 | 0.1 | 0.1 | 0.2 | 0.1 | |
| *Bacteroides uniformis* | 0.516 | | 0.3 | 0.1 | | | | | | 0.2 | | | | | | | | | 0.1 | | | |
| *Anaeromassilibacillus* sp An250 | 0.700 | | | | 0.4 | | | | | 0.1 | | | | | | | 0.2 | | 0.1 | | | |
| *Roseburia inulinivorans* | 0.550 | 0.1 | 0.1 | | | | | | | 0.1 | | | 0.1 | | | | | | 0.1 | | | |
| *Bacteroides caccae* | 0.516 | | 0.1 | | 0.2 | | | | | 0.1 | | | | | | 0.1 | | | | | | |
| *Bacteroides ovatus* | 0.696 | | 0.2 | 0.1 | | | | | | 0.1 | | | | | | | | | | | | |
| *Blautia* sp CAG 257 | 0.665 | | | 0.3 | | | | | | | | 0.9 | | | 0.1 | | | | | | | 0.6 |
| *Blautia hydrogenotrophica* | 0.516 | | | 0.2 | | 0.6 | | | | | | 0.1 | | | 0.3 | | | 0.2 | 0.1 | | | |
| *Denitrobacterium detoxificans* | 0.673 | | | | | | | | | | 0.1 | 0.1 | | | | | | | 0.1 | 0.2 | | |
| *Sellimonas intestinalis* | 0.516 | | | | | 0.4 | | | | | | | | | | | | 0.6 | 0.6 | | | |
| *Bacteroides vulgatus* | 0.516 | | | | 0.1 | | | | | | | | | | | 0.1 | | | 0.1 | | 0.1 | |
| *Eubacterium sp CAG 180* | 0.682 | | | | | | | | | | 0.1 | | | | 0.5 | | 0.3 | | | | | |
| *[Eubacterium] siraeum* | 0.245 | | 0.1 | | | | | | | | | | | | | | 0.1 | | 0.3 | | | |
| *[Clostridium] leptum* | 0.550 | | | 0.1 | 0.2 | | | | | | | | | | | | 0.1 | | 0.1 | | | |

| | | | | | | | | | | | | | | | | | | | | | | |
|---|---|---|---|---|---|---|---|---|---|---|---|---|---|---|---|---|---|---|---|---|---|---|
| *Eisenbergiella tayi* | 0.516 | | | | | | | | | | | | | | | 0.3 | | | | | | |
| *Cloacibacillus evryensis* | 0.516 | | | | | | | | | | | | | | | 0.3 | | | | | | |
| *Blautia hansenii* | 0.516 | | | | | 0.6 | | | | | | | | | | | | | | | 0.2 | |
| *Leuconostoc mesenteroides* | 0.245 | | | | | | | | | | | | 0.4 | | | | | | | | | |
| *Eubacterium* sp CAG 38 | 0.682 | | | | | | | | | | | | 0.1 | | | | | | | | | |
| *Eubacterium ventriosum* | 0.550 | | | | | | | | | | | | | | | 0.1 | | | | | 0.4 | |
| *Ruminococcus* sp CAG 488 | 0.492 | | | | | | | | | | | | | | | 0.4 | | | | | | |
| *Ruminococcus champanellensis* | 0.516 | | | | | | | | | | | | 0.2 | | | 0.1 | | | | | | |
| *Lactonifactor longoviformis* | 0.516 | | | | | 0.2 | | | | | | | | | | | | | | | | |
| *Blautia producta* | 0.516 | | | | | 0.2 | | | | | | | | | | | | 0.1 | | | | |
| *Streptococcus thermophilus* | 0.245 | | | | | | | | | | | | | 0.1 | | | | 0.1 | | | | |
| *Eubacterium* sp CAG OM08 24 | 0.682 | 0.1 | | | | | | | | | | | | | | | | | | | 0.1 | |
| *Eubacterium* sp CAG 274 | 0.682 | 0.1 | 1.0 | | | | | | | | | | | | | | | | | | | |
| *Bacteroides thetaiotaomicron* | 0.696 | | 0.2 | | | | | | | | | | | | | | | | | | | |
| *Clostridium* sp CAG 58 | 0.244 | | | | | | | | | | | | | | | | | | | | | 0.3 |
| | | | | | | | | | | **Indole (IND)** | | | | | | | | | | | | |
| *[Ruminococcus] torques* | 0.292 | 2.0 | 0.8 | 3.3 | | 0.8 | 0.2 | | | 0.9 | 1.0 | 1.6 | 0.2 | 0.2 | 0.7 | 4.2 | 1.8 | 1.5 | 0.1 | 2.2 | 0.0 | 0.9 |
| *Anaerostipes hadrus* | 0.267 | 0.5 | 0.1 | 1.0 | 0.3 | | 0.1 | | 0.7 | | 4.2 | 0.5 | 0.3 | 0.3 | 0.4 | 0.3 | 0.2 | 2.1 | 1.2 | 0.1 | 0.9 | 4.5 |
| *Escherichia coli* | 0.952 | 0.1 | | | 0.1 | | 0.1 | | | 0.2 | | | | | 5.7 | 0.1 | 3.9 | | | | | |
| *Faecalibacterium prausnitzii* | 0.124 | 0.1 | 4.0 | | 0.3 | | | 0.2 | 0.3 | 0.2 | 0.2 | | | | 0.4 | 0.6 | 0.1 | | 0.4 | | 0.1 | 0.1 |
| *Ruminococcus bicirculans* | 0.579 | 0.0 | 1.5 | | 0.6 | | | | | 0.1 | 0.5 | | 0.0 | | | | | | | | | |
| *Bacteroides dorei* | 0.570 | | 1.2 | | | | | | | | | | | | 0.2 | | 0.1 | | | | | 0.0 |
| *Clostridium sporogenes* | 0.570 | | | | | | | | | | | | | 2.9 | | | | | | 0.3 | | |
| *Eubacterium* sp CAG 251 | 0.180 | 1.5 | | | | | | | | | 0.3 | | 0.1 | | 0.2 | | | | | | | |
| *Bacteroides fragilis* | 0.884 | | 0.1 | 0.3 | | | | | | 0.8 | | | | | 1.2 | | | | 0.1 | | | 0.0 |
| *Bacteroides uniformis* | 0.570 | | 0.3 | 0.2 | | | | | | 0.2 | | | | | | | | | 0.2 | | | |
| *Bacteroides ovatus* | 0.765 | | 0.2 | 0.1 | | | | | | 0.1 | | | | | | | | | | | | |
| *Eubacterium ventriosum* | 0.579 | | 0.1 | | | | | | | | | | | | | 0.1 | | | | | 0.4 | |
| *Bacteroides caccae* | 0.570 | | 0.1 | | 0.2 | | | | | 0.1 | | | | | | 0.1 | | | | | | |
| *Alistipes putredinis* | 0.570 | | 0.1 | 0.1 | | | | | | | | | | | | 0.1 | | | 0.1 | | | |
| *Bacteroides vulgatus* | 0.570 | | | | 0.1 | | | | | | | | | | | 0.1 | | | 0.2 | | 0.1 | |
| *Clostridium botulinum* | 0.350 | | | | | | | | | | | | | 0.4 | | | | | | 0.1 | | |
| *Pediococcus_acidilactici* | 0.570 | | | | | | | | | | | | | 0.4 | | | | | | | | |
| *Blautia* sp CAG 257 | 0.116 | | | | | | | | | | | 0.2 | | | | | | | | | | 0.1 |
| *Eubacterium* sp CAG 274 | 0.180 | | 0.3 | | | | | | | | | | | | | | | | | | | |
| *Bacteroides thetaiotaomicron* | 0.765 | | 0.2 | | | | | | | | | | | | | | | | | | | |
| *Clostridium* sp CAG 58 | 0.180 | | | 0.1 | | | | | | | | | | | | | | | | | | 0.6 |
| *Bacteroides cellulosilyticus* | 0.765 | | 0.1 | 0.1 | | | | | | | | | | | | | | | | | | |
| *Actinobaculum* sp oral taxon 183 | 0.843 | | | | | 0.1 | | | | | | | | | | | | | | | | |
| *Eubacterium* sp CAG 180 | 0.180 | | | | | | | | | | | | | | 0.1 | | 0.1 | | | | | |
| *Enterococcus faecalis* | 0.925 | | | | | | | | | | | | 0.1 | | | | | | | | | |
| *Bacteroides_xylanisolvens* | 0.844 | | 0.1 | | | | | | | | | | | | | | 0.1 | | | | | |

| | | | | | | | | | | | | | | | | | | | | | | |
|---|---|---|---|---|---|---|---|---|---|---|---|---|---|---|---|---|---|---|---|---|---|---|
| *Eubacterium limosum* | 0.765 | | | | | | | | | | | | 0.1 | | | | | | | | | |
| *Bacteroides coprophilus* | 0.570 | | | | | | | | | | 0.1 | | | | | | | | | | | |
| *Alistipes finegoldii* | 0.570 | | 0.1 | | | | | | | | | | | | | | | | | | | |
| **Indole lactic acid (ILA)** | | | | | | | | | | | | | | | | | | | | | | |
| *Bifidobacterium longum* | 0.175 | 2.8 | 0.1 | 0.4 | 4.9 | | 1.9 | 1.0 | 2.9 | 8.3 | 1.2 | | 1.9 | 0.6 | 0.3 | 0.2 | 0.1 | 0.3 | 1.1 | 4.3 | | 2.6 |
| *Faecalibacterium prausnitzii* | 0.517 | 0.5 | 16.7 | 0.2 | 1.4 | | 0.1 | 0.7 | 1.4 | 0.6 | 0.7 | 0.1 | 0.1 | | 1.5 | 2.7 | 0.3 | | 1.5 | | 0.4 | 0.3 |
| *Escherichia coli* | 0.676 | 0.1 | | | 0.1 | | | | | 0.1 | | | | | 4.0 | | 2.8 | | | | | |
| *Anaerostipes hadrus* | 0.267 | 0.5 | 0.1 | 1.0 | 0.3 | | 0.1 | | 0.7 | | 4.2 | 0.5 | 0.3 | 0.3 | 0.4 | 0.3 | 0.2 | 2.1 | 1.2 | 0.1 | 0.9 | 4.5 |
| *Bifidobacterium bifidum* | 0.453 | 2.0 | | | 0.3 | | 8.0 | 8.8 | | | | | | 0.8 | | 1.0 | | | | | | |
| *[Eubacterium] rectale* | 0.265 | 2.6 | 0.4 | 1.6 | 0.3 | | | 0.1 | 1.7 | | 0.1 | 0.1 | | | 0.0 | 1.0 | | | 0.4 | | | 0.2 |
| *Bacteroides fragilis* | 0.453 | | | 0.1 | | | | | | 0.4 | | | | | 0.6 | | | | 0.1 | | | |
| *Intestinibacter bartlettii* | 0.124 | | | | | 0.1 | 0.1 | | | | | | 0.2 | | | | | | 0.1 | | | |
| *Bifidobacterium adolescentis* | 0.265 | | 0.1 | 0.1 | | | | | | 0.1 | | | | | | | | | 0.1 | | | |
| *Bacteroides ovatus* | 0.265 | | 0.1 | | | | | | | | | | | | | | | | | | | |
| *Bacteroides thetaiotaomicron* | 0.265 | | 0.1 | | | | | | | | | | | | | | | | | | | |
| **Skatole (SK)** | | | | | | | | | | | | | | | | | | | | | | |
| *[Eubacterium] rectale* | 0.598 | 5.8 | 0.9 | 3.7 | 0.6 | | | 0.1 | 3.9 | 0.1 | 0.2 | 0.2 | | | | 2.3 | | 0.1 | 0.8 | | | 0.38 |
| *Intestinibacter bartlettii* | 0.380 | 0.1 | | | | 0.2 | 0.3 | | | 0.1 | 0.1 | | 0.6 | 0.1 | | | | 0.1 | 0.2 | | | 0.05 |
| *Parabacteroides distasonis* | 0.380 | | | 0.1 | | | | | | | | | | | | | | | | | | |
| **Indole propionic acid (IPA)** | | | | | | | | | | | | | | | | | | | | | | |
| *Clostridium sporogenes* | 0.350 | | | | | | | | | | | | | 1.8 | | | | | | 0.2 | | |
| *Clostridium botulinum* | 0.867 | | | | | | | | | | | | | 0.9 | | | | | | 0.2 | | |
| *Clostridium_paraputrificum* | 0.350 | | | | | | 0.1 | | | | | | | | | | | | | | | |
| **Indole acrylic acid (IA)** | | | | | | | | | | | | | | | | | | | | | | |
| *Clostridium sporogenes* | 0.480 | | | | | | | | | | | | | 2.5 | | | | | | 0.3 | | |
| **Indole (carbox)aldehyde (I-Ald)** | | | | | | | | | | | | | | | | | | | | | | |
| *[Ruminococcus] gnavus* | 0.803 | 0.5 | | 0.9 | | 2.1 | | | | | | | | | 0.3 | | | | 0.1 | | | 1.0 |
